# The genomic epidemiology of multi-drug resistant nontyphoidal *Salmonella* causing invasive disease in sub-Saharan Africa

**DOI:** 10.1101/2020.12.11.422246

**Authors:** Se Eun Park, Duy Thanh Pham, Gi Deok Pak, Ursula Panzner, Ligia Maria Cruz Espinoza, Vera von Kalckreuth, Justin Im, Ondari D. Mogeni, Heidi Schütt-Gerowitt, John A. Crump, Robert F. Breiman, Yaw Adu-Sarkodie, Ellis Owusu-Dabo, Raphaёl Rakotozandrindrainy, Abdramane Bassiahi Soura, Abraham Aseffa, Nagla Gasmelseed, Arvinda Sooka, Karen H. Keddy, Jürgen May, Peter Aaby, Holly M. Biggs, Julian T. Hertz, Joel M. Montgomery, Leonard Cosmas, Beatrice Olack, Barry Fields, Nimako Sarpong, Tsiriniaina Jean Luco Razafindrabe, Tiana Mirana Raminosoa, Leon Parfait Kabore, Emmanuel Sampo, Mekonnen Teferi, Biruk Yeshitela, Muna Ahmed El Tayeb, Ralf Krumkamp, Denise Myriam Dekker, Anna Jaeger, Adama Tall, Aissatou Niang, Morten Bjerregaard-Andersen, Sandra Valborg Løfberg, Jessica Fung Deerin, Jin Kyung Park, Frank Konings, Sandra Van Puyvelde, Mohammad Ali, John D. Clemens, Gordon Dougan, Stephen Baker, Florian Marks

## Abstract

**Background:** Invasive nontyphoidal *Salmonella* (iNTS) is one of the leading causes of bacteraemia in sub-Saharan Africa. Multi-drug resistance (MDR) and further resistance to third generation cephalosporins and fluoroquinolones have emerged in multiple iNTS serotypes. Molecular epidemiological investigations of nontyphoidal *Salmonella* are needed to better understand the genetic characteristics and transmission dynamics associated with major MDR iNTS serotypes across the continent.

**Methods:** A total of 166 nontyphoidal *Salmonella* isolates causing invasive disease were collected from a multi-centre study in eight African countries between 2010 and 2014, and whole-genome sequenced to investigate the geographical distribution, antimicrobial genetic determinants and population structure of iNTS serotypes-genotypes. Phylogeographical reconstruction was further conducted in context of the existing genomic framework of iNTS serotypes Typhimurium and Enteritidis. Population-based incidence of MDR-iNTS disease was also estimated.

**Results:** *Salmonella enterica* subsp. *Enterica* serotype Typhimurium (*S*. Typhimurium) sequence-type (ST) 313 and *Salmonella enterica* subsp. *Enterica* serotype Enteritidis (*S*. Enteritidis) ST11 were predominant, and both exhibited high frequencies of MDR. *Salmonella enterica* subsp. *Enterica* serotype Dublin (*S*. Dublin) ST10 emerged in West Africa. Mutations in the *gyrA* gene were identified in *S*. Enteritidis and *S*. Typhimurium in Ghana; and ST313 carrying *bla*_CTX-M-15_ was found in Kenya. Inter-country transmission of MDR ST313 lineage II and the West African Clade of MDR ST11 between Ghana and neighbouring countries including Mali, Burkina Faso, and Nigeria were evident. The incidence of MDR-iNTS disease exceeded 100/100,000-person years-of-observation (PYO) in children aged <5 years in several West African countries.

**Conclusions:** Multiple MDR iNTS serotypes-sequence types, predominantly *S*. Typhimurium ST313 and *S*. Enteritidis ST11, are co-circulating in sub-Saharan Africa with evidence of transmission between West African countries. The development of safe and effective iNTS vaccines coupled with appropriate antimicrobial stewardship and adequate epidemiological monitoring are essential to limit the impact of these pathogens in Africa.

## Background

Non-typhoidal *Salmonella enterica* are classically considered a collective of zoonotic pathogens associated with self-limiting diarrhoea in humans. However, certain nontyphoidal *Salmonella* serovars have become an established cause of invasive disease (iNTS) in specific geographical regions, particularly affecting immunocompromised individuals including infants and young adults with HIV, malaria and malnutrition [1]. Annually, there are an estimated 3.4 million cases of iNTS disease globally, 20% of these are fatal [2],[3]. The vast majority of iNTS diseases occur in sub-Saharan African countries where annual incidences up to 175-388 per 100,000 person-years and case fatality ratio (CFR) as high as 25% have been reported in children aged <5 years [4],[5],[6],[7],[8]. A recent multicentre study across sub-Saharan Africa also reported iNTS as a major cause of bacteraemia in febrile patients in Africa, with adjusted incidences >100 per 100,000 person-years identified across multiple sampling locations [9].

Various *Salmonella* serovars can cause iNTS disease; however, previous studies have shown that the majority of iNTS infections in sub-Saharan Africa are caused by two predominant serovars, namely *Salmonella enterica* subsp. *Enterica* serotype Typhimurium (*S*. Typhimurium) and *Salmonella enterica* subsp. *Enterica* serotype Enteritidis (*S*. Enteritidis) [10],[11],[12],[13]. *S*. Typhimurium in sub-Saharan Africa are largely associated with a single multi-drug resistant (MDR) genotype, known as ST313, comprised of two major lineages I and II [14]. *S*. Enteritidis are estimated to account for a third of the iNTS burden in sub-Saharan Africa and are primarily associated with the ST11 genotype. Further, there are three major clades of *S*. Enteritidis ST11 (Global epidemic, West African and Central/East African clades) co-circulating in this region with a rapid increase in MDR [15]. In addition to *S*. Typhimurium and *S*. Enteritidis, other *Salmonella enterica* subsp. *Enterica* serovars associated with iNTS disease include *S*. Isangi [16], *S*. Concord [17], *S*. Stanleyville, and *S*. Dublin [18], signifying the complex epidemiological landscape of iNTS in sub-Saharan Africa.

Historically, ampicillin, chloramphenicol, and trimethoprim-sulfamethoxazole (co-trimoxazole) have been the first-line drugs for the treatments of iNTS disease as well as typhoid fever in Africa [19]. The increased use of antimicrobials has consequently led to the emergence and widespread of MDR, defined as resistant to these first-line drugs, in iNTS organisms; particularly *S*. Typhimurium and *S*. Enteritidis [10],[20]. Numerous sub-Saharan African countries have reported high prevalences of MDR iNTS disease, including Malawi [20-22], Kenya [23],[24],[25], Ghana [26], Gambia [5], Democratic Republic of the Congo (DRC) [27], Mozambique [7,28], Tanzania [29], Burkina Faso [9], Guinea-Bissau [9], and Nigeria [30]. Consequently, alternative treatments, such as ciprofloxacin, azithromycin, and ceftriaxone, are being increasingly deployed to manage such infections. However, these drugs are unavailable or costly in some resource-limited settings [13],[22]. Their effectiveness is likely diminished further due to the emergence of MDR plus ceftriaxone resistance as well as extensively drug-resistance (XDR), defined as MDR combined with resistance to fluoroquinolones (i.e. ciprofloxacin) and third generation cephalosporins (i.e. ceftriaxone), as reported in *S*. Typhimurium ST313 in Kenya [19], Malawi [12], and the DRC [31]. The continental dissemination of MDR organisms together with the emergence of highly resistant variants pose a significant challenge for the treatment, management, and control of iNTS disease in Africa [19].

Here, through whole genome sequencing (WGS), we aimed to investigate the distribution of iNTS serovars and their corresponding sequence types (ST) and antimicrobial resistance (AMR) determinants through a contemporaneous collection of iNTS strains from multiple sites across sub-Saharan Africa. We additionally estimated the incidence of MDR iNTS disease in the sampling locations and performed comprehensive phylogeographical analyses of *S*. Typhimurium ST313 and *S*. Enteritidis ST11 in the context of globally representative collection of respective organisms.

## Methods

### Study design and inclusion criteria

Between 2010 and 2014, standardized healthcare facility-based surveillance of invasive *Salmonella* infections in sub-Saharan African populations was conducted in thirteen sites in ten countries (Typhoid Fever Surveillance in Africa Program, TSAP); the methods were published [32]. In brief, febrile patients from all age groups (except in Ghana, where only children aged <15 years were enrolled) with a tympanic or axillary temperature of ≥38.0°C or ≥37.5 °C, respectively, living in a defined study catchment area were eligible for recruitment. For inpatients, reported fever within the period of 72 hours prior to admission was added to the inclusion criteria. Written informed consent/assent was obtained. Clinical assessments of patients included history and current of illness, physical examination, and clinical appraisal. Eligible patients were treated per the respective national guidelines. Blood samples (5-10 mL for adults; 1-3 mL for children) were collected for laboratory diagnostics of these febrile cases.

### Bacterial isolates and antimicrobial susceptibility testing

Through the TSAP study, blood specimens were obtained from febrile patients; one aerobic blood culture bottle per patient. Collected bottles were incubated in systems with automated growth detection (BACTEC Peds Plus Medium/BACTECT Plus Aerobic-F, BACTEC, Becton-Dickinson, New Jersey; or BacT/ALERT PF Paediatric FAN/BacT/ALERT FA FAN Aerobic, bioMerieux, Marcy l’Etoile, France) except for the study site in Sudan whereby manual blood culture techniques were used. Blood cultures with bacterial growth were sub-cultured on solid media such as blood agar and chocolate agar (Oxoid, Basingstoke, United Kingdom), and biochemical tests were conducted (API 20E, bioMerieux) to confirm suspected *Salmonella* isolates [32]. Confirmatory testing was conducted at the study reference laboratories at Oxford University Clinical Research Unit (OUCRU, Ho Chi Minh city) and Bernhard Nocht Institute for Tropical Medicine (BNITM) using the VITEK automate. Antimicrobial susceptibility testing was performed using agar diffusion tests according to the Clinical Laboratory and Standards Institute (CLSI) guidelines [32]. Malaria diagnostics were performed via a combination of thick and thin blood smears and rapid malaria test (SD Bioline Malaria Ag P.f./P.v., SD Standard Diagnostics, Suwon, Republic of Korea), this was variable between sites [33].

### Data sources and bacterial isolates

The total of 166 iNTS isolates was used for this investigation, which comprised 94 iNTS isolates from the TSAP study, an additional 23 iNTS isolates detected outside the predefined study catchment area, and 49 iNTS isolates from other studies conducted in Agogo, Ghana in 2007-2009. For further analyses on the *S*. Typhimurium and *S*. Enteritidis isolates in context of the global phylogeny of respective organisms, existing datasets were incorporated: 102 iNTS serovar Typhimurium ST313 Lineage II isolates from seven countries (Malawi, Kenya, Mozambique, Uganda, DRC, Nigeria, and Mali) [10], Nigeria and DRC [11], Malawi [12], Kenya [13], Malawi [34], and 594 iNTS serovar Enteritidis ST11 isolates (selected from Feasey *et al*. 2016) [15]. A summary of these isolates and their country of origin is provided in Supplementary Table 1.

### Whole genome sequencing

Genomic DNA was extracted from all *Salmonella* isolates using the Wizard Genomic DNA Extraction Kit (Promega, Wisconsin, USA). Two μg of genomic DNA from each organism was subjected to indexed-tagged pair-end sequencing on an Illumina Hiseq 2000 platform (Illumina, CA, USA) to generate 100 bp paired-end reads. Raw sequence data are available in the European Nucleotide Archive (Project number: ERP009684, ERP010763, ERP013866).

### Single Nucleotide Polymorphism (SNP) calling and analyses

Raw Illumina reads were used to create multiple assemblies using Velvet v1.2 [35] with parameters optimized using VelvetOptimiser v2.2.5 [36],[37] and automated annotation was performed using PROKKA v1.5 [38]. Roary [39] was used to define the pan genome of 166 iNTS isolates with blastp percentage identity of 99% and a core definition of 99%. 3,450 core genes were identified (genes that present in ≥99% strains) and 86,765 SNP sites were extracted from the core gene alignment using SNP-sites [37].

For *S*. Typhimurium ST313, raw Illumina reads of 99 isolates from this study and additional 102 isolates of *S*. Typhimurium ST313 lineage II from previous studies [10],[11],[12],[13],[34] were mapped to the reference sequence of *S*. Typhimurium strain SL1344 (accession: FQ312003.1), using SMALT version 0.7.4 (http://www.sanger.ac.uk/resources/software/smalt/). Candidate SNPs were called against the reference sequence using SAMtools [40] and filtered with a minimum mapping quality of 30 and minimum consensus base agreement of 75%. The allele at each locus in each isolate was determined by reference to the consensus base in that genome using SAMtools *mpileup* and removing low confidence alleles with consensus base quality ≤20, read depth ≤5 or a heterozygous base call. Repeat finding program in NUCmer v3.1 [41] was used to identify exact repetitive regions of ≥ 20 bp in length in the reference genome and SNPs called in these regions were excluded. SNPs called in phage sequences and recombinant regions identified using Gubbins [42] were further removed, resulting in a final set of 1,960 chromosomal SNPs. The identification of SNPs for *S*. Enteritidis ST11 was performed following the same procedure as *S*. Typhimurium ST313. Briefly, the raw Illumina reads of 28 *S*. Enteritidis ST11 isolates from my studies and additional 594 isolates from a global collection [15] were mapped to the reference sequence of *S*. Enteritidis strain P125109 (accession: NC_011294.1), using SMALT followed by SNP calling and filtering as above resulting in a final set of 25,121 SNPs.

### Phylogenetic analyses

A maximum likelihood (ML) phylogenetic tree was constructed from the 86,765 SNP alignment of all 166 iNTS isolates using RAxML version 8.2.8 with a generalized time-reversible model and a Gamma distribution to model the site-specific rate variation (GTRGAMMA in RAxML) [43], and outgroup rooted. Clade support for this tree was assessed through a bootstrap analysis with 100 pseudo-replicates. To investigate the molecular epidemiology of my *S*. Typhimurium ST313 and *S*. Enteritidis ST11 isolates in regional and international context, ML tree was inferred from an alignment of 1,960 SNPs for 201 *S*. Typhimurium ST313 lineage II isolates (99 from this study and 102 from previous studies [10],[11],[12],[13],[34] and an alignment of 25,121 SNPs for 622 *S*. Enteritidis ST11 isolates (28 from this study and 594 from a global collection [15], using RAxML with the same parameters as above. Support for these phylogenetic trees was assessed through a 100 bootstrap pseudo-analysis. Tree annotation was visualized using ITOL [44].

### Antimicrobial resistant gene and plasmid analyses

From raw Illumina reads, Short Read Sequence Typer-SRST2 [45] was used to identify the acquired AMR genes and their precise alleles using the ARG-Annot database [46], as well as the plasmid replicons using the PlasmidFinder database [47]. Multilocus sequence typing (MLST) of all iNTS isolates was also determined using SRST2 together with MLST database for *Salmonella enterica* downloaded from pubMLST. *Salmonella* serovars were identified using conventional serology as well as MLST [48] and genomics [49]. To investigate the isolates with resistant genotypes, a *de novo* genome assembly was performed from raw sequences using SPAdes [50] version 3.11.0, followed by Bandage [51] to inspect the location of AMR genes in the corresponding assembled contigs.

### Incidence analyses of MDR iNTS disease

Incidence of MDR iNTS was estimated per 100,000-person years-of-observation (PYO) for MDR iNTS isolates detected from Burkina Faso, Ghana, Guinea-Bissau, Kenya, and Senegal. Statistical methodology used previously to calculate the incidence of *S*. Typhi and iNTS disease in TSAP countries was used to calculate MDR iNTS incidence [9],[32],[52]. Briefly, an age-stratified PYO were estimated using available demographic data in HDSS (Health and

Demographic Surveillance System) and non-HDSS sites and health-seeking behaviour of randomly selected individuals, representative of the study population, were factored in (denominator). The recruitment proportion was adjusted to the age-stratified crude MDR iNTS cases (numerator). Adjusted incidence of MDR iNTS per 100,000 PYO was estimated with 95% CIs using these adjustment factors and crude MDR iNTS case numbers. The previously established multi-country database (FoxPro software) for TSAP was used for the countries with MDR iNTS isolates.

## Results

### The geographical distribution of iNTS serovars and sequence types in Africa

The majority of iNTS infections in sampled sub-Saharan African countries were caused by two predominant serovars-STs, *S*. Typhimurium ST313 and *S*. Enteritidis ST11; however, their distribution varied between countries (Table 1). More specifically, 66% (110/166) of the isolates were *S*. Typhimurium, of which 90% (99/110) were ST313 and 10% (11/110) ST19. *S*. Enteritidis accounted for 18% (30/166) of the isolates, comprising ST11 (93.3%; 28/30), ST183 (3.3%; 1/30) and ST2107 (3.3%; 1/30). Notably, *S*. Dublin (ST10) also accounted for 11% (18/166) of the iNTS isolates. We also found several less common serovars-STs including *S*. Choleraesuis ST145 (3/166), *S*. Muenster ST321 (1/166), *S*. Poona ST308 (1/166), *S*. Stanleyville ST339 (1/166), and *S*. Virchow ST359 (1/166).

**Table 1.** The distribution of iNTS serovars and genotypes circulating in the sampled countries in sub-Saharan Africa

| Country<br>(no. of iNTS<sup>1</sup>)<br>(a) | Serovars | No. of iNTS<br>per serovar<br>(b) | % of total no.<br>of iNTS per<br>country<br>(b)/(a) | Genotype<br>(Sequence<br>type) | No. of<br>iNTS<br>per<br>Sequence<br>type<br>(c) | % of total<br>no. of iNTS<br>per country<br>(c)/(a) |
| --- | --- | --- | --- | --- | --- | --- |
| Burkina Faso (12) | Typhimurium | 7 | 58% | 313 | 6 | 50% |
|  |  |  |  | 19 | 1 | 8% |
|  | Enteritidis | 4 | 33% | 11 | 4 | 33% |
|  | Dublin | 1 | 8% | 10 | 1 | 8% |
| Ghana (133) | Typhimurium | 92 | 69% | 313 | 90 | 68% |
|  |  |  |  | 19 | 2 | 2% |
|  | Enteritidis | 20 | 15% | 11 | 18 | 14% |
|  |  |  |  | 183 | 1 | 1% |
|  |  |  |  | 2107 | 1 | 1% |
|  | Dublin | 17 | 13% | 10 | 17 | 13% |
|  | Muenster | 1 | 1% | 321 | 1 | 1% |
|  | Poona | 1 | 1% | 308 | 1 | 1% |
|  | Stanleyville | 1 | 1% | 339 | 1 | 1% |
|  | Virchow | 1 | 1% | 359 | 1 | 1% |
| Guinea Bissau (9) | Typhimurium | 5 | 56% | 313 | 2 | 22% |
|  |  |  |  | 19 | 3 | 33% |
|  | Enteritidis | 1 | 11% | 11 | 1 | 11% |
|  | Choleraesuis | 3 | 33% | 145 | 3 | 33% |
| Kenya (1) | Typhimurium | 1 | 100% | 313 | 1 | 100% |
| Madagascar (4) | Typhimurium | 1 | 25% | 19 | 1 | 25% |
|  | Enteritidis | 3 | 75% | 11 | 3 | 75% |
| Senegal (2) | Typhimurium | 1 | 50% | 19 | 1 | 50% |
|  | Enteritidis | 1 | 50% | 11 | 1 | 50% |
| South Africa (1) | Enteritidis | 1 | 100% | 11 | 1 | 100% |
| Tanzania (4) | Typhimurium | 3 | 75% | 19 | 3 | 75% |
|  | Unknown | 1 | 25% | 2533 | 1 | 25% |
| Total 8<br>Countries<sup>2</sup><br>(166 iNTS<br>isolates)(d) | Serovars | No. of iNTS<br>per serovar<br>(e) | % of total no.<br>of iNTS in all<br>8 countries<br>(e)/(d) | Genotype<br>(Sequence type) | No. of iNTS<br>per<br>genotype<br>(f) | % of total no.<br>of iNTS in all<br>8 countries<br>(f)/(d) |
|  | Typhimurium | 110 | 66% | ST313 | 99 | 60% |
|  |  |  |  | ST19 | 11 | 7% |
|  | Enteritidis | 30 | 18% | ST11 | 28 | 17% |
|  |  |  |  | other STs<sup>3</sup> | 2 | 1% |
|  | Dublin | 18 | 11% | ST10 | 18 | 11% |
|  | Others | 8 | 5% | Other STs<sup>4</sup> | 8 | 5% |

<sup>1</sup> iNTS: invasive nontyphoidal *Salmonella*
<sup>2</sup> Total 8 countries: Burkina Faso, Ghana, Guinea-Bissau, Kenya, Madagascar, Senegal, South Africa, and Tanzania presented in this table. No iNTS isolates were yielded in Sudan and Ethiopia.
<sup>3</sup> Other sequence types of *S. Enteritidis* detected: 1 ST183 (isolate yielded from Ghana; age/sex unknown due to missing data) and 1 ST2107 (from Ghana; a 22 year old female); both non-MDR and no antimicrobial resistant genes detected.
<sup>4</sup> Sequence types of the other NTS serovars: 1 Muenster ST321 (yielded from Ghana; age/sex/year unknown due to missing data), 1 Poona ST308 (yielded from Ghana in 2008; age/sex unknown due to missing data), 1 Stanleyville ST339 (yielded from Ghana; age/sex/year unknown due to missing data), 1 Virchow ST359 (from Ghana; age/sex/year unknown due to missing data), 3 Choleraesui ST145 (2 isolates yielded from 1 year old female infants in 2010 and 2011 and 1 from 3 year old female infant in 2012; all from Guinea-Bissau), 1 Unknown ST2533 from Tanzania.

Regarding the geographical distribution of iNTS, *S*. Typhimurium ST313 were predominately found in West Africa (Burkina Faso, Ghana, Guinea-Bissau). In East and Southern Africa, ST313 was exclusively detected in Kenya (Table 1). *S*. Enteritidis ST11, though less common than ST313, appeared to be pervasive in both West (Burkina Faso, Ghana, Guinea-Bissau, Senegal) and Southern (South Africa, Madagascar) Africa. Additionally, *S*. Dublin ST10 was only detected in Burkina Faso and Ghana in West Africa. *S*. Typhimurium ST19 appeared geographically dispersed across the continent, including West (Burkina Faso, Ghana, Guinea-Bissau, Senegal) and East and Southern Africa (Tanzania, Madagascar).

Overall, 61% (102/166) of the iNTS organisms were multi-drug resistant. MDR iNTS organisms were isolated in Burkina Faso (83%; 10/12), Ghana (66%, 88/133), Guinea-Bissau (22%; 2/9), Kenya (100%; 1/1), and Senegal (50%; 1/2). *S*. Typhimurium exhibited the highest prevalence of MDR (85%; 94/110), and notably 95% (94/99) of the ST313 isolates were MDR. 23% (7/30) of *S*. Enteritidis and 6% (1/18) *S*. Dublin were MDR. None of the 11 *S*. Typhimurium ST19 isolates were MDR (Figure 1, Table 2).

**Figure 1.**
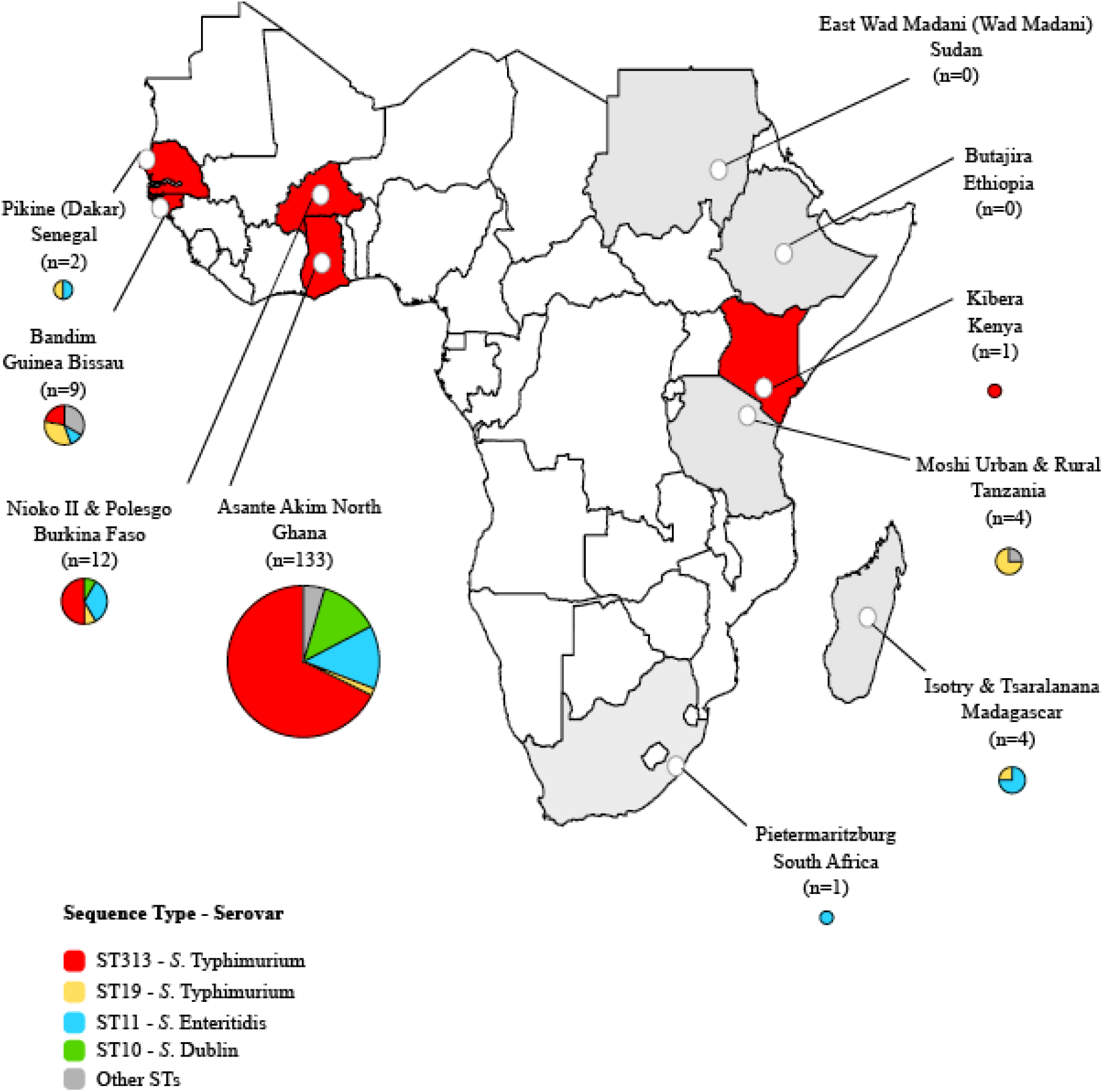
The geographical distribution of iNTS genotypes and serovars in the sampled countries in sub-Saharan Africa. Different colours in the pie charts correspond to different sequence types and serovars of iNTS isolates in our study sites. The size of the pie charts corresponds to the numbers of isolates in each country. Countries highlighted in red and grey are with and without MDR iNTS isolates respectively.

**Table 2.** The distribution of MDR iNTS and *gyrA* mutation (fluoroquinolone resistance) in the sampled countries in sub-Saharan Africa

| **Serovars**<br>**(number of isolates)**<br>**N=166** | **MDR iNTS**<br>**isolates per**<br>**serovars** | **MDR iNTS serovar**<br>**per country**<br>**(%, number)** | **MDR iNTS**<br>**per genotype**<br>**(%, number)** |
| --- | --- | --- | --- |
| Typhimurium (n=110) | 85% (94/110) | Burkina Faso (50%, 6/12)<br>Ghana (64%, 85/133)<br>Guinea Bissau (22%, 2/9)<br>Kenya (100%, 1/1) | ST313 (95%, 94/99)<br>ST19 (0%, 0/11) |
| Enteritidis (n=30) | 23% (7/30) | Burkina Faso (33%, 4/12)<br>Ghana (2%, 2/133)<br>Senegal (50%, 1/2) | ST11 (25%, 7/28) |
| Dublin (n=18) | 6% (1/18) | Ghana (1%, 1/133) | ST10 (6%, 1/18) |
| Others (n=8) | 0% (0/8) | n.a. | n.a. |

| **Countries**<br>**(no. of iNTS)** | **No. of MDR**<br>**iNTS (N=102)** | **% of MDR**<br>**iNTS** | ***gyrA*** |
| --- | --- | --- | --- |
| Burkina Faso (n=12) | 10 | 10/12 (83%) | 0 |
| Ghana (n=133) | 88 | 88/133 (66%) | S83Y (ST313, 2 isolates)<br>D87G (ST11, 6 isolates)<br>D87Y (ST11, 2 isolates)<br>D87N (ST11, 3 isolates) |
| Guinea-Bissau (n=9) | 2 | 2/9 (22%) | 0 |
| Kenya (n=1) | 1 | 1/1 (100%) | 0 |
| Senegal (n=2) | 1 | 1/2 (50%) | 0 |

<sup>1</sup> Multidrug resistance (MDR) definition used for the analysis presence of resistant genes for at least one agent in all three antimicrobial categories listed below (detected in this study): Ampicillin (blaCTX\_M, blaOXA, blaTEM), Chloramphenicol: (*catA1*), Trimethoprim-sulfamethoxazole (sulfonamide (sul1, sul2) and trimethoprim (*dfrA*, *dfrA1*, *dfrA14*, *dfrA8*)).
<sup>2</sup> Refer to Table 1 for the number of iNTS isolates per country used as a denominator to calculate the % of MDR per country in this table.
<sup>3</sup> The 2 MDR iNTS isolates with *gyrA* mutation (fluoroquinolone resistance) were yielded from a 1 year old female infant and 10 month old female infant in Agogo in 2011 (TSAP: Typhoid Fever Surveillance Program).
<sup>4</sup> *Spv* locus were detected in all MDR iNTS isolates.

### Phylogenetics and AMR determinants of iNTS serovars

Our phylogenetic reconstruction of all iNTS isolates showed that the three major serovars-STs: *S*. Typhimurium (ST313), *S*. Enteritidis (ST11), and *S*. Dublin (ST10) formed three distinct clusters and exhibited dissimilar AMR gene profiles (Figure 2). Almost all of the MDR *S*. Typhimurium ST313 (95%; 94/99) carried the Tn21 transposon-associated MDR-loci (*sulII-strAB-dfrA1-aadA1-sulI-cat-blaTEM*) on an IncF virulent-resistance plasmid pSLT-BT as reported before [10]. One MDR *S*. Typhimurium ST313 from Kenya additionally carried two copies of *bla*_CTX-M-15_, conferring resistance to third generation cephalosporins, one of which was located on the 300kb IncHI2 plasmid, pKST313 (accession number: LN794248). The other *bla*_CTX-M-15_ was inserted into the chromosome and disrupted the *ompD* locus. Of 99 *S*. Typhimurium ST313, 5 (5.1%) harboured pSLT-BT, but were not associated with MDR due to the lack of MDR-*loci*. Reduced suseptibility to fluoroquinolones was uncommon in *S*. Typhimurium ST313, with 2% (2/99) of Ghanaian ST313 isolates possessing a mutation (S87Y) in *gyrA* (Table 2).

**Figure 2.**
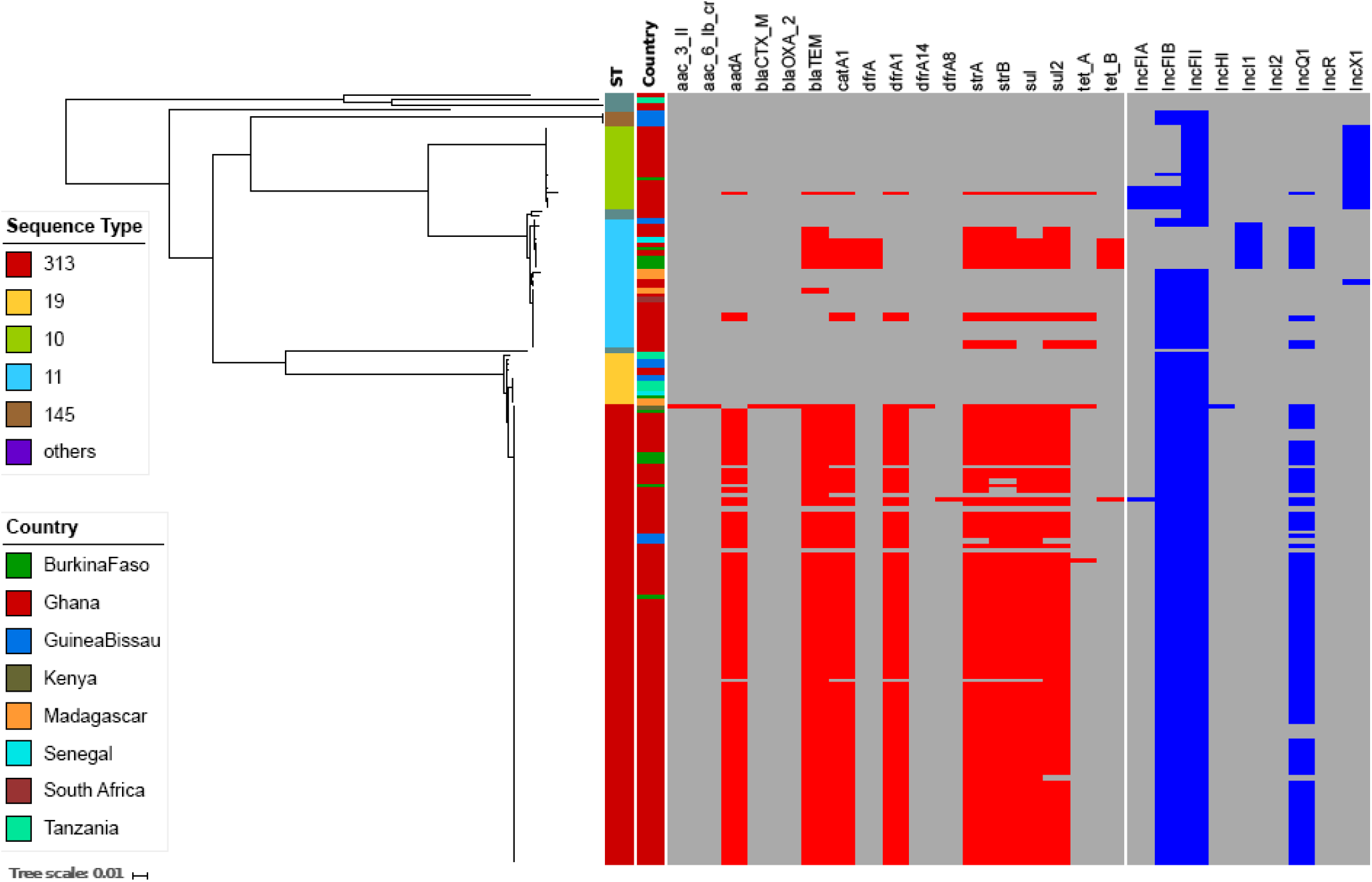
Antimicrobial resistant genes and plasmids associated with iNTS isolates circulating in sampled sub-Saharan Africa countries. Maximum likelihood phylogenetic tree based on the core genes of all our iNTS isolates and their corresponding metadata. First column shows the sequence types in different colour. Second column corresponds to the countries where our iNTS isolates were detected. The remaining columns exhibit a heatmap of detected AMR genes and plasmid replicons. The tree scale bar indicates the number of substitutions per variable site.

The majority of *S*. Enteritidis ST11 (19/28; 68%) harboured a typical IncF virulence plasmid (60kb), which was comparable to pSENV (accession number NC_019120.1, coverage 100%, identity 99%). The remaining *S*. Enteritidis ST11 (9/28; 32%) harboured a novel IncI1 virulence-resistance plasmid (pSEP, accession number: ERP121368) of approximately 68kb (Figure 3), of which 7/28 (25%) isolates (4/Burkina Faso, 2/Ghana, 1/Senegal) carried the MDR-encoding Tn21-like transposon (*sulII-strAB-tetB-dfrA1-sulI-cat-blaTEM*), and 2/28 (7.1%) isolates carried an AMR cassette (*TnpA-sulII-strAB-blaTEM-Tn3*), conferring resistance against ampicillin, streptomycin, and sulphanomides. The novel IncI plasmid exhibited 60% homology to pSENV and did not harbour the IncF replicons, the *pefBACD* fimbriae-encoding operon, or the virulent-associated genes *srgA* and *rck* (Figure 3). In addition, two non-MDR Ghanaian *S*. Enteritidis ST11 possessed an AMR cassette (*sul2-strAB-tetA*) associated with a small (11kb) non-conjugative IncQ plasmid conferring resistance against sulphonamides, streptomycin and tetracyclines. This IncQ plasmid exhibited a similar genetic structure to pSTU288-2 from *S*. Typhimurium (accession number CP004059.1, coverage 98%, identity 99%). Two further non-MDR Ghanaian isolates carried a Tn21-mediated AMR cassette (*sulII-strAB-dfrA1-aadA1-sulI-cat*) on the virulence plasmid, and a single non-MDR isolate from Madagascar carried a *bla*TEM-bearing Tn3 integrated into the virulence plasmid. Reduced suseptibility to fluoroquinolones was predicted in 39% (11/28) of the *S*. Enteritidis-ST11, all of which originated from Ghana, displaying differing mutations in codon 87 in *gyrA* (D87G: 6 isolates, D87N: 3 isolates, and D87Y: 2 isolates) (Table 2).

**Figure 3.**
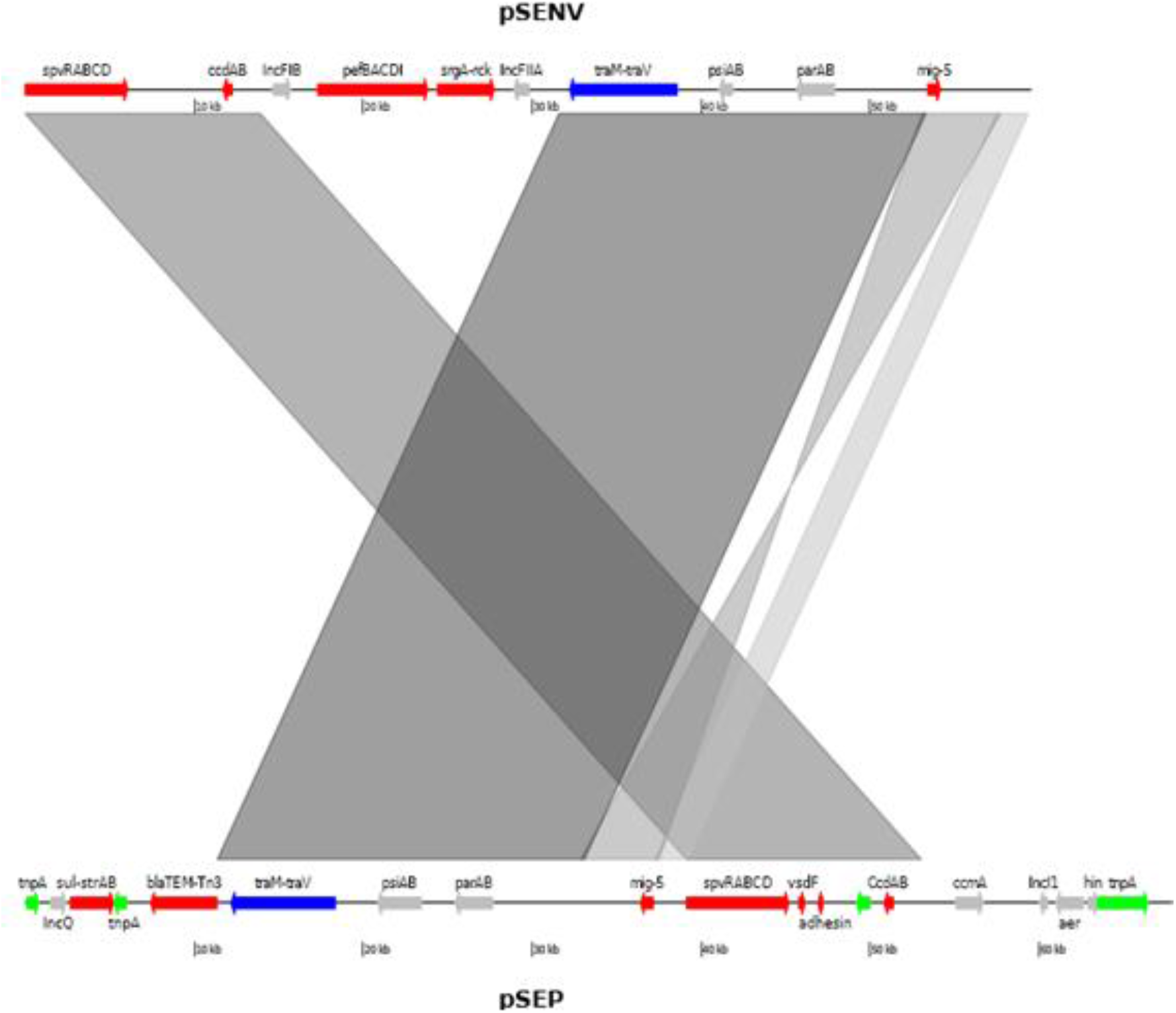
Novel IncI1 virulence-resistance plasmid (pSEP) in an *S*. Enteritidis ST11 isolate. Plasmid comparison analyses between the novel virulence-resistance IncI1 plasmid pSEP (bottom) and the reference virulence IncF plasmid pSENV (top). The grey blocks show the BLASTn comparison between the two plasmids. Some annotations are added for both plasmids. Red coloured arrows are genes associated with virulence and AMR. Blue coloured arrows are genes associated with conjugation. Grey coloured arrows correspond to plasmid replication and stability. Green coloured genes are associated with transposon elements.

Most *S*. Dublin ST10 isolates (17/18; 94%) did not carry any AMR genes, except for one Ghanaian isolate with an MDR-encoding Tn21 transposon-mediated cassette (*sulII-strAB-dfrA1-aadA1-sulI-cat-blaTEM)* (Figure 2). All *S*. Dublin ST10 isolates carried a typical IncF virulence plasmid (75kb), similar to pOU1115 (accession number: NC_010422.1, coverage 100%, identity 99%). None of the *S*. Dublin ST10 isolates possessed mutations in *gyrA* (Table 2). Markedly, the *viaB* operon encoding the Vi polysaccharide was absent from all *S*. Dublin isolates. Lastly, all *S*. Typhimurium ST19, as well as other uncommon serovars-STs, had no acquired AMR genes and considered pan-antimicrobial susceptible.

### Phylogeography of S. Typhimurium ST313 and S. Enteritidis ST11 isolates in global context

To investigate the potential transmission patterns of predominant iNTS serovars of *S*. Typhimurium ST313 and *S*. Enteritidis ST11 in a broader context, we reconstructed the phylogenetics of our isolates together with previously published data. We found that all *S*. Typhimurium ST313 isolated from our study belong to lineage II, suggesting lineage I may no longer be circulating in the sampled countries. One Kenyan *S*. Typhimurium ST313 with two copies of *bla*_CTX-M-15_ formed part of the previously described clonal expansion of the MDR ceftriaxone-resistant ST313 sublineage in Kenya [19]. The ST313 from Ghana displayed a high degree of genetic diversity and scattered around the phylogenetic tree. This observation indicates these organisms have been likely circulating in Ghana for a prolonged period. The ST313 from Ghana clustered with isolates from Mali, Burkina Faso, and Nigeria on several occasions, indicating frequent movement of organisms between Ghana and the neighbouring West African countries (Figure 4a).

**Figure 4.**
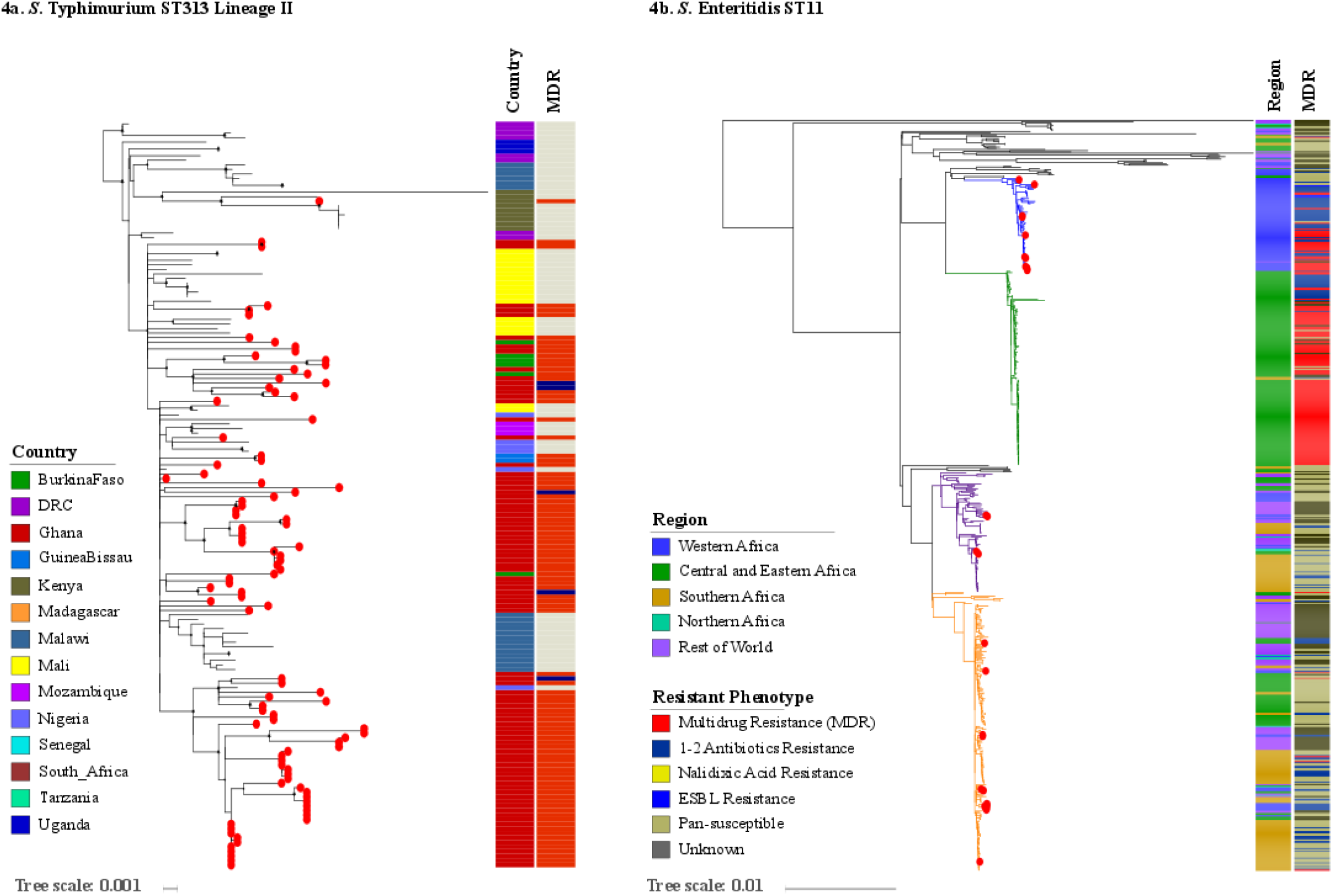
The phylogeography of *S*. Typhimurium ST313 lineage II and *S*. Enteritidis ST11 in sub-Saharan Africa. **a)** Maximum likelihood phylogenetic tree of my *S*. Typhimurium-ST313 lineage II isolates in the context of sub-Saharan Africa continent. Red circles at the terminal leaves correspond to our study isolates. First column shows different colour-coded countries from where all analysed isolates originate. Second column shows MDR and non-MDR isolates in red and grey respectively. **b)** Maximum likelihood phylogenetic tree of my *S*. Enteritidis-ST11 isolates in the global context of *S*. Enteritidis. Red circles at the terminal leaves correspond to our study isolates. First and second columns show regions and resistant phenotypes in different colours. The tree scales indicate the number of substitutions per variable site.

Further phylogeographical investigation of *S*. Enteritidis ST11 in a global context demonstrated that, 11/28 (40%) (Ghana: 5, Burkina Faso: 4, Senegal: 1, Guinea Bissau: 1) isolates fell into the West African clade described by Feasey et al, known to be associated with AMR and an invasive disease phenotype[15]. Our ST11 organisms within this West African clade also displayed an MDR (7 isolates) or AMR phenotype (2 isolates). Within this clade, we found evidence that some Ghanaian isolates clustered tightly with isolates from Burkina Faso and previously described isolates from Mali. Additionally, 13/28 (46%) of the *S*. Enteritidis ST11 (Ghana 11, Madagascar: 1, South Africa: 1) belonged to the Global epidemic clade described by existing publication, of which these isolates appeared as part of their country-specific endemic circulation; however, we also found that some Ghanaian isolates clustered with previously described isolates from its neighbouring Cameroon and Senegal. We identified 4/28 (14%) of the ST11 isolates grouped within the Outlier cluster, of which 2 Ghanaian isolates and two isolates from Madagascar clustered with previously described isolates from the U.S. and Mauritius, respectively. All these organisms were susceptible to all antimicrobials (Figure 4b).

### Incidence of MDR iNTS disease in sub-Saharan Africa

The age stratified incidences of MDR iNTS in Burkina Faso, Ghana, Guinea-Bissau, and Kenya are illustrated in Table 3. The adjusted incidence of MDR iNTS disease exceeded 100/100,000-person years-of-observation (PYO) in children <15 years of age in all West African countries: Burkina Faso (NiokoII, 274/100,000 PYO, 95% confidence interval [CI], 185-406; and Polesgo, 255/100,000 PYO, 95% CI, 138-470), Ghana (Asante Akim North: Ghana-AAN, 414/100,000, 95% CI, 333-515), and Guinea-Bissau (Bandim, 105/100,000, 95% CI, 69-161). Among children <15 years, younger children (<2-4 years) exhibited the highest MDR iNTS incidences in both sites in Burkina Faso: 753/100,000 PYO (95% CI, 460-1233) in NiokoII and 630/100,000 PYO (95% CI, 288-1380) in Polesgo. In both settings in Burkina Faso, the incidence of MDR iNTS disease in the infant age-group was slightly lower than in the 2-4-year olds though all children <5 years exhibited a high burden of MDR iNTS disease. In Ghana-AAN, infants aged <2 year had the highest incidence of MDR iNTS disease (1435/100,000; 95% CI, 1110-1854) followed by children aged between 2 and 5 years (747/100,000; 95% CI, 491-1135). Similarly in Guinea-Bissau, infants <2 years exhibited the highest incidence of MDR iNTS disease (291/100,000; 95% CI, 176-482). The incidence rate of MDR iNTS in older age groups (≥15 years) was relatively low, ranging between 0 (Guinea-Bissau), to 11 (Kenya), and 54 (Burkina Faso) per 100,000 PYO.

**Table 3.**
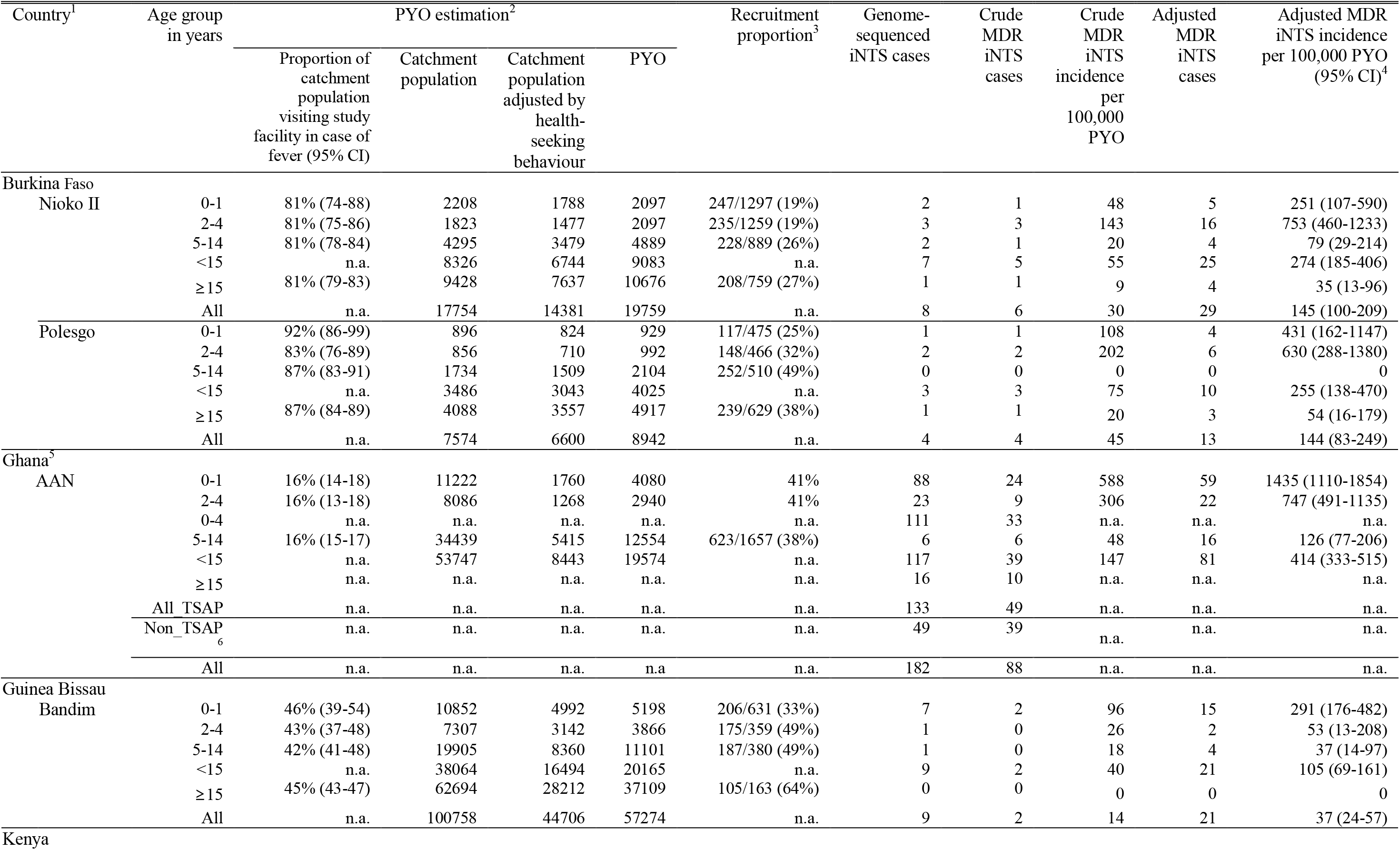
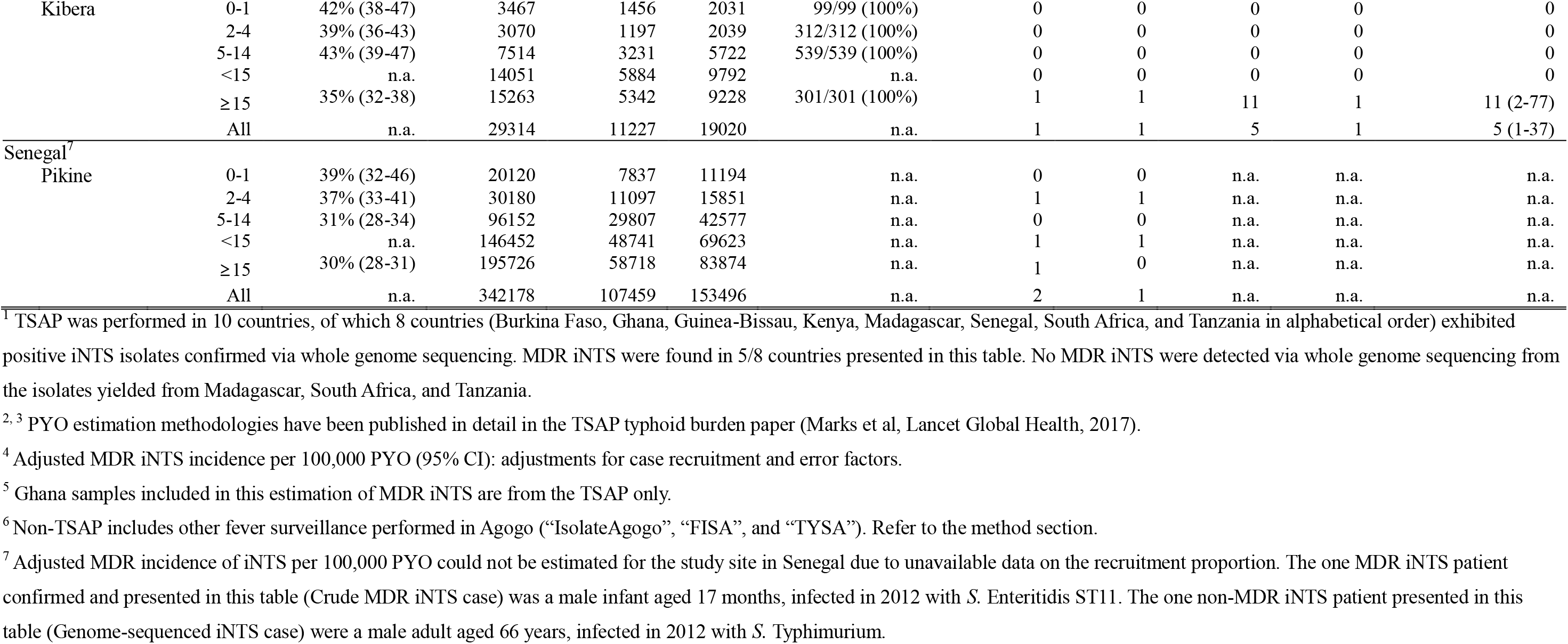
Incidence estimates of MDR iNTS disease in sub-Saharan Africa

## Discussion

Although much understanding about iNTS disease in Africa has been gained over the last decade, the disease remains a major public health problem in parts of sub-Saharan Africa. While there is currently no vaccine for iNTS disease, the disease burden is further exacerbated by the regional dissemination of MDR phenotypes and the emergence of resistance against newer drugs in some countries. Moreover, iNTS disease has been caused by a large number of *Salmonella* serovars with indistinguishable clinical symptoms [2]. NTS serovars may exhibit different propensity to develop drug resistance and have dissimilar host range and transmission patterns [13],[53]. It is therefore critical to monitor the distribution of iNTS serovars and their AMR patterns over time and across the continent to facilitate updated treatment guidelines for local public health authorities and healthcare workers.

Our study found high frequency of MDR iNTS infections in the sampled countries in sub-Saharan Africa. Despite the fact that MDR *S*. Typhimurium ST313 remains the most common cause of iNTS disease, our data highlight the emergence of other less common MDR iNTS serovars; particularly MDR *S*. Enteritidis ST11 and *S*. Dublin ST10 in West Africa. Recent studies have also reported that MDR *S*. Enteritidis ST11 appears to be an emerging cause of bloodstream infections in Africa [54],[55]. The source and predominant modes of transmission for these increasingly important iNTS serovars are currently unknown and requires further investigation. In addition, the co-circulation of multiple MDR iNTS serovars poses a substantial challenge for the control and prevention measures. Vaccines for iNTS disease need to target multiple key *Salmonella* serotypes and such efforts are currently underway including a bivalent O-antigen polysaccharide-CRM197 conjugate vaccine targeting both *S*. Typhimurium and *S*. Enteritidis, using Generalized Modules for Membrane Antigens (GMMA) technology [56] and an iNTS vaccine program by the International Vaccine Institute (IVI) [57].

Typically, iNTS serovars possess a non-conjugative virulence plasmid of IncF replicon type; as a result, the acquisition of the novel virulence-resistance IncI1 plasmid carrying both the *spv* locus and MDR-encoding cassette in *S*. Enteritidis ST11 is worrying as such plasmid may be further disseminated via horizontal gene transfer. This phenomenon also raises the possibility that different iNTS serovars may have access to different resistance gene pools and may have capacities to increase virulence and/or resistance. Routine surveillance with the integration of WGS should be implemented to closely monitor the trend in antimicrobial resistance and dominant iNTS serovars as well as to timely detect novel AMR clones that are epidemiologically and clinically important for informing public health and clinical practices.

Our reconstructed phylogeography of the two predominant MDR serovars-genotypes (*S*. Typhimurium ST313 and *S*. Enteritidis ST11) indicated all contemporaneous *S*. Typhimurium ST313 fell into lineage II with evidence of occasional transfer between Ghana and the neighbouring countries (Burkina Faso, Guinea-Bissau, Senegal, and Nigeria) in West Africa. The ST11 isolates belonged to three distinct lineages (Global epidemic clade, West African clade and an Outlier cluster), of which all MDR ST11 isolates fell within the West Africa clade and appeared to be widely distributed throughout Ghana, Burkina Faso, and Senegal. The observation that multiple ST11 lineages are found to cause bloodstream infections is worrying as these organisms are associated with multiple transmission pathways and may have the ability to acquire additional virulence-resistance plasmids [58]. Previous studies suggest that asymptomatic carriers are likely an important reservoir of nontyphoidal *Salmonella* [59],[60], which may contribute to the transmission of these pathogens within and between countries. Given the complex transmission mechanisms of iNTS/NTS serovars, one health genomic approach should be also considered in an attempt to better understand the sources and transmission routes of various iNTS/NTS serovars across different ecological niches. Further, understanding population mobility across the continent will contribute to monitoring disease expansion and the success of local efforts at control.

Antimicrobial therapy is essential to treat patients with iNTS disease and to reduce mortality. However, the effectiveness of antimicrobial therapy is diminishing due to the emergence and dissemination of various MDR and XDR NTS serovars and strains [15],[31],[61]. Our data provides further insights into the emergence of reduced susceptibility to fluoroquinolones among MDR ST313 and ST11 from Ghana as well as the continued circulation of ceftriaxone-resistant ST313 sublineage in Kenya. Invasive *Salmonella* with reduced susceptibility to ciprofloxacin have also been increasingly reported in Burkina Faso [9], Ghana [9,26], Nigeria [62], Senegal [63], Mozambique [28], the Congo [64], and South Africa [65]. This increasing trend in AMR against clinically important classes of newer antimicrobials among multiple iNTS serovars in Africa is of particular concern given the fact that patients infected with highly resistant organisms are likely not to respond to treatment with these newer drugs [19]. It is probable that the widespread use of ceftriaxone and ciprofloxacin for the treatment of febrile illnesses in Africa will eventually lead to a rapid increase and spread of MDR and XDR pathogens in this continent, which may amplify the burden of the disease.

We also found a substantial incidence of MDR iNTS disease in the West African countries including Burkina Faso, Ghana (<5 years), and Guinea-Bissau (< 2 years). The incidence of MDR iNTS disease presented here generally corresponded to the incidence of iNTS disease in respective countries [9]. Among these Western African countries, Ghana in particular exhibited high incidences of both MDR iNTS disease and MDR typhoid fever reported previously [66], indicative of concurrence of iNTS disease and typhoid fever in Ghana with increasing public health concern associated with invasive MDR salmonellosis. Conversely, in East African countries such as Kenya and Tanzania, MDR iNTS was less prominent relative to the high incidence of MDR typhoid fever in some of the same locations [66].

Our study did not take into account the prior usage of antimicrobials nor other clinical features reported with iNTS disease such as gastrointestinal symptoms [67], and our enrolment criteria excluded afebrile patients and those unable to provide informed consent. As a result, our incidence calculation may be an under-estimation of the true burden of MDR iNTS disease among the study populations. Different sample sizes across sampled locations in respective countries may limit a direct comparison on AMR occurrence between regions, and thus adjusted incidence estimations may be more suitable for data comparability. The substantial burden of MDR iNTS disease in infants and young children in these countries is likely to undermine the effectiveness of therapy, which may lead to prolonged illness and increased risks to case fatalities. For instance, a hospital-based study of iNTS disease in Malawian children exhibited high prevalence of MDR iNTS disease and identified significant risk factors for mortality, including a history of dyspnea, absence of fever, presence of respiratory distress or hypoglycaemia at presentation, and HIV infection [67]. Further studies on MDR and iNTS disease severity are warranted.

## Conclusions

We have found high prevalence of MDR iNTS infections in sub-Saharan African countries which are predominantly caused by *S*. Typhimurium ST313 and *S*. Enteritidis ST11. Our study also provides further insights into the transmissions of MDR ST313 lineage II and the West African Clade of MDR ST11 between West African countries. High incidence of MDR-iNTS disease in children aged <5 years is found in several West African countries. Antimicrobial stewardship and continuous surveillance and investigations on the severity and fatalities associated with iNTS disease are instrumental to guide appropriate treatment therapy and enhance control and prevention measures. In addition, the deployment of a safe and effective polyvalent vaccine against the most commonly found iNTS serotypes should be prioritized for the management and prevention of iNTS disease in Africa, particularly in countries where there is high prevalence of MDR iNTS infections, malaria and malnutrition.

### Declarations

#### Ethics approval and consent to participate

This research was conducted under the ethical principles of the Declaration of Helsinki. The IVI Institutional Review Board (IRB), the national ethical review committees in each participating country, and local research ethics committees approved the study protocol. All eligible patients meeting the study inclusion criteria were provided with a detailed explanation of the study purpose, and written informed consent was obtained prior to study enrolment. For children, written informed consent was obtained from parent or guardian [68].

## Supporting information

The genomic epidemiology of multi-drug resistant nontyphoidal Salmonella causing invasive disease in sub-Saharan Africa

## Consent for publication

Not applicable.

## Availability of data and materials

Raw sequence data are available in the European Nucleotide Archive (projects ERP009684, ERP010763, ERP013866). SMALT version 0.7.4 used is available at: http://www.sanger.ac.uk/resources/software/smalt/.

## Competing interests

The authors declare no competing interests.

## Funding

This study was supported by the Bill & Melinda Gates Foundation (grant: OPPGH5231). The findings and conclusions are our own and do not necessarily reflect positions of the Bill & Melinda Gates Foundation or the US Centers for Disease Control and Prevention. The International Vaccine Institute acknowledges its donors, including the Government of Republic of Korea and the Swedish International Development Cooperation Agency (SIDA). Research infrastructure at the Moshi site was supported by the US National Institutes of Health (R01TW009237; U01 AI062563; R24 TW007988; D43 PA-03-018; U01 AI069484; U01 AI067854; P30 AI064518), and by the UK Biotechnology and Biological Sciences Research Council (BB/J010367). SB is a Sir Henry Dale Fellow, jointly funded by the Wellcome Trust and the Royal Society (100087/Z/12/Z). DTP is funded as a leadership fellow through the Oak Foundation (OCAY-15-547).

## Author’s contributions

SEP conducted the phylogenetic, resistance gene and plasmid analyses and interpretation of results under the scientific supervision of DTP. SEP and GDP analysed incidence estimations. SEP structured, drafted, and edited the paper under the scientific guidance from SB and DTP. FM, SEP, VvK, LMCE, UP, GDP, JI, OM, HSG, JAC, RFB, YAS, EOD, RR, ABS, AA, NG, KHK, JM, AGS, PA, HMB, JTH, JMM, LC, BO, BF, NS, TJLR, TMR, LPK, ES, MT, BY, MAET, AS, RK, DMD, AJ, AT, AN, MBA, SVL, JFD, JKP, and FK contributed to data acquisition in the field, interpretation of results, and editing of the paper. All authors read and approved the final draft.

## Acknowledgements

We would like to acknowledge and thank all staff and partners involved in obtaining and processing the data and samples including healthcare facility and laboratory staff at the TSAP network countries. We also thank the WTSI Pathogen Informatics team for help with whole genome sequencing.

## Notes

### Competing Interest Statement

The authors have declared no competing interest.

