## Supplementary material for "The genomic epidemiology of multi-drug resistant nontyphoidal *Salmonella* causing invasive disease in sub-Saharan Africa": The genomic epidemiology of multi-drug resistant nontyphoidal Salmonella causing invasive disease in sub-Saharan Africa

### Supplementary Table 1. All iNTS organisms analysed

| | | | | | | | | |
|---|---|---|---|---|---|---|---|---|
| | **All iNTS isolates from TSAP[2] & non-TSAP projects[3] in Ghana (N=166)** | | | | | | | |
| **iNTS isolates analyzed[1] (N=945)** | Burkina Faso | Ghana[4] | Guinea-Bissau | Kenya | Madagascar | Senegal | South Africa | Tanzania |
| | 12 | 133 | 9 | 1 | 4 | 2 | 1 | 4 |
| | 4 (Polesgo) | 84 (Asante Akim North) | 6 (Simao Hospital) | 1 (Kibera) | 3 (CHU Tsaralalana) | 1 (IPS) | | 2 (Moshi Urban) |
| | 8 (NiokoII) | 49 (Agogo)[5] | 2 (Bandim) | | 1 (Isotry) | 1 (Dominique hospital) | | 1 (Moshi Rural) |
| | | 36 (“IsolateAgogo”) | 1 (Belem) | | | | | 1 (Moshi Other) |
| | | 5 (“FISA”) | | | | | | |
| | | 3 (“TYSA”) | | | | | | |
| | | 5 (unidentified) | | | | | | |
| | **Published iNTS Typhimurium ST313 Lineage II isolates (N=102)[6]** | | | | | | | |
| | DRC[7] | Kenya | Malawi | Mali | Mozambique | Nigeria | Uganda | |
| | 8 | 9 | 55 | 18 | 3 | 6 | 3 | |
| | **Published iNTS Enteritidis ST11 isolates (N=677)[8]** | | | | | | | |
| | Western Africa | Central & Eastern Africa | Southern Africa | Northern Africa | Rest of the World | | | |
| | 90 | 262 | 131 | 11 | 183 | | | |

[1] Total 945 iNTS organisms were analysed in this manuscript: 166 iNTS isolates from the TSAP and other Ghana projects; sequenced and published 102 iNTS serovar typhimurium ST313 Lineage II isolates from seven countries (DRC/Kenya/Malawi/Mali/Mozambique/Nigeria/Uganda); and sequenced and published 594 (out of 677) iNTS serovar Enteritidis ST11 isolates (See appendices for the metadata).

[2] TSAP: Typhoid Fever Surveillance Program (Von Kalckreuth *et al.* 2016; Marks *et al.* 2017). Of the ten countries under the TSAP, no iNTS was found in Sudan and Ethiopia. 4 and 1 iNTS strains were identified from Madagascar and South Africa, respectively, through the whole genome sequencing; no iNTS isolates were reported from Madagascar and South Africa based on the blood culture result. Few analysed iNTS isolates, which may have been detected outside the strictly predefined surveillance catchment area, are considered for the bacterial genomic analysis, but excluded from the incidence estimation of the multidrug resistant iNTS in respective sites in Table 4. The 166 iNTS isolates included for the final genomic analyses are based on the screening of whole genome sequenced results.

[3] Non-TSAP projects in Ghana include the “Febrile Illnesses Surveillance in Africa (FISA)”, “Typhoid Surveillance in Africa (TYSA)”, and “IsolateAgogo” conducted in Agogo.

[4] Ghana: 133 iNTS isolates analysed in this manuscript include 84 iNTS detected through the TSAP in Asante Akim North and 49 iNTS from several non-TSAP projects conducted in Agogo.

[5] 49 (Agogo): 49 iNTS isolates detected from various non-TSAP projects in Agogo, Ghana as listed in this table.

[6] Total 102 iNTS Typhimurium ST313 Lineage II isolates, which had been collected in seven countries in sub-Saharan Africa and whole genome sequenced, were published and raw sequence data accessible at the Wellcome Trust Sanger Institute. Global collection of *S.* Typhimurium and additional *S.* Typhimurium from: Malawi, Kenya, Mozambique, Uganda, DRC, Nigeria, and Mali (Okoro *et al.* 2012); Nigeria and DRC (Leekitcharoenphon *et al.* 2013); Malawi (Feasey *et al.* 2014); Kenya (Kariuki *et al.* 2015b); Malawi (Msefula *et al.* 2012). Whole genome sequenced 166 iNTS isolates from TSAP and non-TSAP projects in Ghana listed in this table included 99 iNTS Typhimurium ST313 Lineage II . This 99 ST313 Lineage II was combined with the raw sequence data of 102 ST313 Lineage II published isolates listed in this footnote. Total 201 ST313 Lineage II isolates were further analysed.

[7] DRC: Democratic Republic of Congo

[8] Raw sequence data of the published global collection of 594/677 *S.* Enteritidis isolated from human and animal samples (Feasey *et al.* 2016) were accessible at the Wellcome Trust Sanger Institute. The associated metadata were publicly available online. These samples were collected and whole genome sequenced across the African continent and the world:

- Western Africa: Benin, Cameroon, Chad, Gabon, Guinea, Ivory Coast, Mali, Niger, and Senegal
- Central & Eastern Africa: Central African Republic, Comoros, Congo, Djibouti, Ethiopia, HPA, Kenya, Madagascar, Malawi, Mauritius, Mozambique, Rwanda, and Uganda
- Southern Africa: Republic of South Africa
- Northern Africa: Algeria, Egypt, Mauritania, Morocco, and Tunisia
- Rest of the World: Angola, Argentina, Columbia, Germany, Italy, Japan, PTG, Ratin-manufactured rat poison, Slovakia, Spain, Thailand, UK, Uruguay, USA, and unknown
